# Inducing oncolytic cell death in human cancer cells by the long non-coding RNA *let-A*

**DOI:** 10.1101/2021.07.16.452707

**Authors:** Tosca Birbaumer, Tommy Beat Schlumpf, Makiko Seimiya, Yanrui Jiang, Renato Paro

## Abstract

Long non-coding (lnc) RNAs contain functional elements that play important regulatory roles in a variety of processes during development, normal physiology, as well as disease. We recently discovered a new lncRNA, we named *let-A*, expressed from the evolutionary conserved *let-7-Complex* locus in *Drosophila*. This RNA induces cell death in *Drosophila* cancer cells. Here we show that ectopic expression of *Drosophila let-A* is also exerting an oncolytic toxicity in several human cancer cell lines, but shows almost no effect in more differentiated or cell lines derived from normal tissue. We demonstrate that *let-A* RNA prepared by *in vitro* transcription and provided in the growth medium is sufficient to induce cell death both in human and *Drosophila* cancer cells. The activity of *in vitro* transcribed *let-A* is most efficient in its full length, but requires prior modification/processing to become active. *let-A* induces a reduction of nucleolar size in treated cells. We show exo/endocytosis and Toll signaling pathway to be necessary for *let-A*-induced toxicity. Our findings indicate *let-A* exhibits an evolutionary conserved anti-cancer function, making it a promising molecule for tumor treatments.

## Introduction

Long non-coding RNAs (lncRNAs) are a class of RNA molecules that are usually transcribed from intergenic and intronic regions. lncRNAs are more than 200 nucleotides long and not translated. Computational sequence analyses have predicted tens of thousands lncRNAs encoded in the genome of an organism. Large-scale transcriptome analyses further confirmed the expression of these RNA species in different cells in temporally highly coordinated patterns. Acting either *in cis* or *in trans*, the functions of only few lncRNAs have been experimentally characterized, including regulation of transcription and splicing, organization of nuclear structures, and control of other RNA molecules or as prosthetic group in protein complexes (Kopp & Mendell, 2018; Ransohoff et al., 2018; Ulitsky & Bartel, 2013). The exact biological functions of the majority of annotated lncRNAs remain elusive, however.

Over the past decade, lncRNAs have also become a focus in cancer research. Pan-cancer genome and transcriptome analyses have revealed hundreds of lncRNAs that are either mutated or differentially expressed, and act as either onco-RNA (*MALAT*, *HOTAIR*) or tumor-suppressive RNA (*LincRNA-p21*) in various types of cancer (Slack & Chinnaiyan, 2019). As described in our accompanying paper (Birbaumer et al., 2021), we identified a new lncRNA transcript named as *let-A* in the *Drosophila let-7-Complex*. *let-A* is encoded in the first intron of the *let-7-C* locus, and is transiently expressed during normal development in response to the molting hormone ecdysone. Induced expression of *let-A* results in rapid cell death of the *in vitro* cultured *Drosophila ph^505^* cancer cells, which was established from *Drosophila ph^505^* mutant imaginal disc cells (Birbaumer et al., 2021). Such conditioned culture medium, or purified RNA molecules from the medium then becomes toxic to other cancer cells. After incubation with *let-A* conditioned medium, *in vivo* grown tumor tissue also becomes non-tumorigenic upon re-transplantation. In addition, feeding such induced medium to host flies carrying transplanted tumors leads to reduced growth of tumors in these animals. In this accompanying paper, we present data showing that *Drosophila let-A* is also able to exert its toxic activity when ectopically expressed in various types of mammalian cancer cell lines. This toxicity was not observed on more differentiated or human cell lines derived from normal tissue. We show that *in vitro* transcribed *let-A* RNA is sufficient to impair both human and *Drosophila* cancer cells, and is most efficient in its full length. In addition, *let-A* RNA prepared *in vitro* requires prior modification/processing with either *Drosophila* or mammalian cell extracts to become active. We show fluorophore-labeled *let-A* can enter cancer cells. Furthermore, cellular exo/endocytosis and Toll signaling pathway are required for *let-A*-induced toxicity.

## Results

### *Drosophila let-A* RNA exhibits its toxic effect also on mammalian cancer cells

As we described in our accompanying paper (Birbaumer et al., 2021), lncRNA *let-A* can induce rapid cell death in *Drosophila* cancer cells *in vitro* as well *in vivo*. We therefore tested if this RNA shows a similar effect on mammalian cells, by transducing HEK293T cells with *Drosophila let-A* and measuring consequences on cellular viability. To our surprise, *Drosophila let-A* was also toxic to HEK293T cells as most cells died overnight after induction (Fig. 1A). Furthermore, the RNA fraction purified from the medium of *ph^505^*cells transduced with *let-A* (conditioned let-A/medium) was also able to eliminate HEK293T cells, whereas RNAs from medium of *I1B* or mCherry-transduced cells used as controls could not (Fig. 1B). These results showed *Drosophila let-A* RNA can exert its surprising toxicity even on mammalian cells.

**Figure 1.**
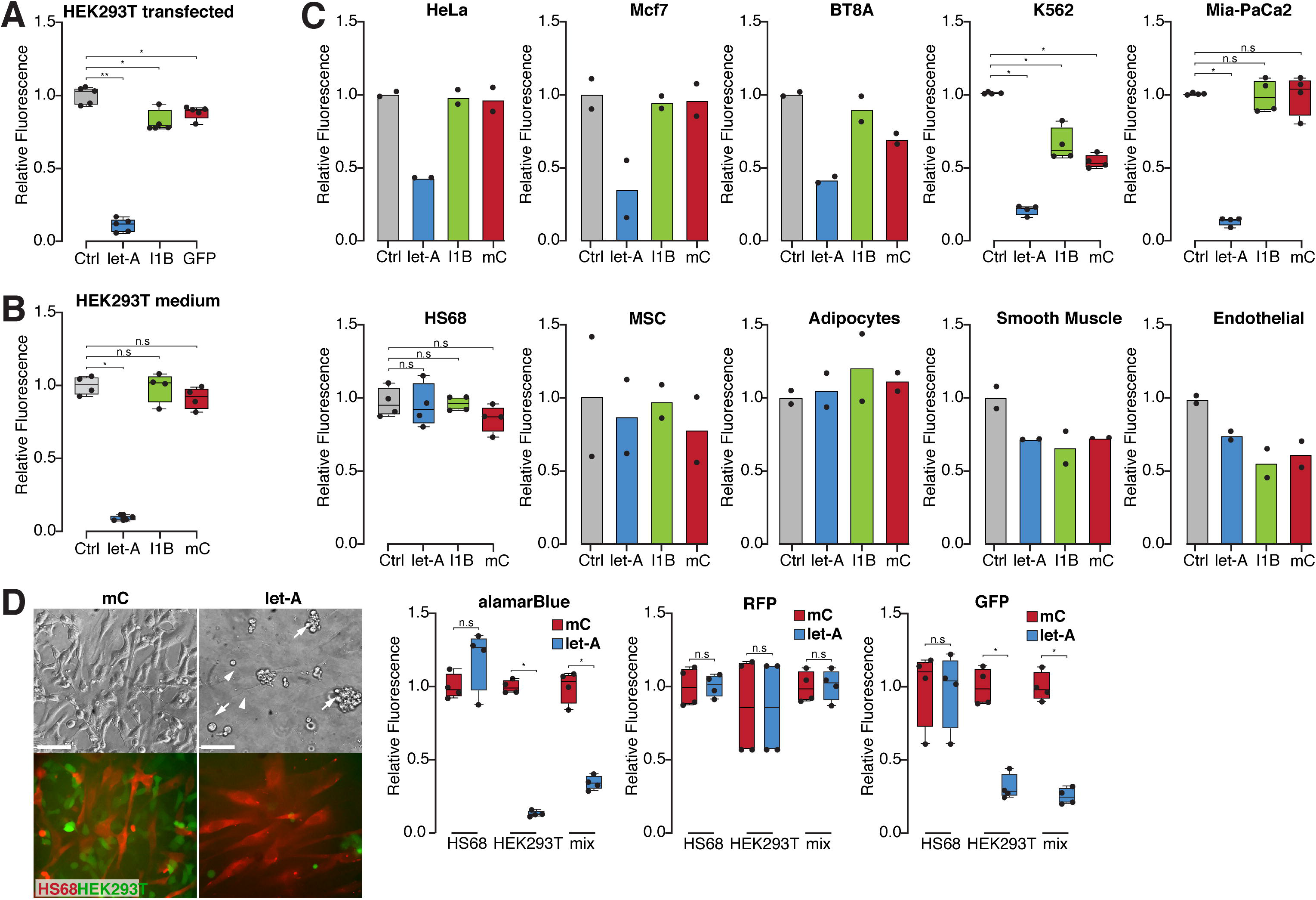
*Drosophila let-A* can induce cell death in mammalian cultured cancer cells. **(A)** HEK293T cells were transfected with constructs expressing the indicated RNA *let-A*, I1B (please refer to (Birbaumer et al., 2021), and GFP. After induction only *let-A* resulted in cell death as measured by alamarBlue. I1B or GFP had only minor effects on cell viability. Bars across conditions denote Wilcoxon Rank-Sum Test p-values (n.s.: non-significant, *: ≤ 0.05, **: ≤ 0.01). **(B)** HEK293T cells were treated with RNA, purified from *let-A*, I1B, or mCherry (mC) conditioned *ph^505^* cell cultures. Only RNA from let-A/medium resulted a comparable reduction of HEK293T cell viability. Conversely, little effect was observed with RNA from I1B/medium or mC/medium. Bars across conditions denote Wilcoxon Rank-Sum Test p-values (n.s.: non-significant, *: ≤ 0.05). **(C)** A number of different mammalian cell lines were treated with RNA purified from *let-A* conditioned medium obtained from *ph^505^* transduced culture. This included HeLa (human cervical cancer), Mcf7 (human breast cancer), BT8A (human glioblastoma), K562 (human leukemia), and Mia PaCa-2 (human pancreatic cancer) cells. Purified RNA from let-A/medium was able to kill all the cell of these cell lines but RNAs purified from I1B/medium or mCherry/medium did not show an effect. Surprisingly, cell lines derived from a non-tumorigenic source did not respond to *let-A*, these include HS68 (human fibroblast cells), MSC (human primary mesenchymal stem cells), and differentiated adipocyte, smooth muscle cells, and endothelial cells. For figures with bars, the bars across conditions denote Wilcoxon Rank-Sum Test p-values (n.s.: non-significant, *: ≤ 0.05) **(D)** When a mixed culture of HEK293T (GFP, green) and HS68 (RFP, red) are treated with let-A/medium, only HEK293T cells undergo rapid cell death while the fibroblast cells survive the treatment, as quantified by alamarBlue and RFP/GFP fluorescence. Bars across conditions denote Wilcoxon Rank-Sum Test p-values (n.s.: non-significant, *: ≤ 0.05).

We next treated a number of different mammalian cell lines with RNA purified from *let-A* conditioned medium obtained from a *ph^505^* transduced culture. These included HeLa (human cervical cancer), Mcf7 (human breast cancer), BT8A (human glioblastoma), K562 (human leukemia), and Mia PaCa-2 (human pancreatic cancer) cells (Fig. 1C). Indeed, purified RNA was able to kill all the cell of these lines (Fig. 1C), but RNAs purified from I1B/medium or mCherry/medium did not show an effect (Fig. 1C). As such, *Drosophila let-A* appears to have an evolutionarily conserved oncolytic function which is also active in mammalian cells. Interestingly, we observed some cell lines that did not respond to *let-A*, including HS68 (human fibroblast cells), MSC (human primary mesenchymal stem cells), as well as differentiated adipocyte, smooth muscle cells, and endothelial cells (Fig. 1C), suggesting that mammalian tumor-derived cells are more prone to *let-A* toxicity compared to cells derived from more differentiated tissue.

To further test the specificity of *let-A*, we co-cultured HEK293T cells and fibroblast-derived HS68 cells on the same plate. Interestingly, overnight incubation of the co-culture with let-A/medium resulted in the elimination of most HEK293T cells, but showed little effect on HS68 cells (Fig. 1D). Taken together, these results suggest that *let-A* exhibits also in a mixed population a specific toxicity towards mammalian tumorigenic/immortal cell lines, but shows less of an effect on cell lines derived from a non-tumorigenic source.

### *In vitro* transcribed *let-A* RNA needs further processing to become active

To clarify if the efficiency of the toxicity is caused by *let-A* RNA, we *in vitro* transcribed (ivt) sense and antisense *let-A* RNA or I1B and mCherry RNA as controls, and tested their toxicity first on *Drosophila ph^505^* cells. When *ph^505^* cells were treated with purified ivt *let-A* RNA, cell viability was not affected. However, when the ivt *let-A* RNA was first incubated with *Drosophila* cell extracts, then added to *ph^505^* cells, cells were killed (Fig. 2A and supplementary figure 1A). Additionally, activated ivt *let-A* RNA shows a similar toxicity to mammalian HEK293T cells (Fig. 2A). Indeed, only RNA with the sense sequence could kill the cells, but RNA with the antisense sequence to *let-A* could not. When ivt *let-A* RNA was sonicated before incubation with extract and applied to *ph^505^* cells, it was no longer able to kill *ph^505^* cells (Fig. 2B). These results indicate the effector molecule to be *let-A*, however, requiring further processing and/or modifications to induce the cell toxicity.

**Figure 2.**
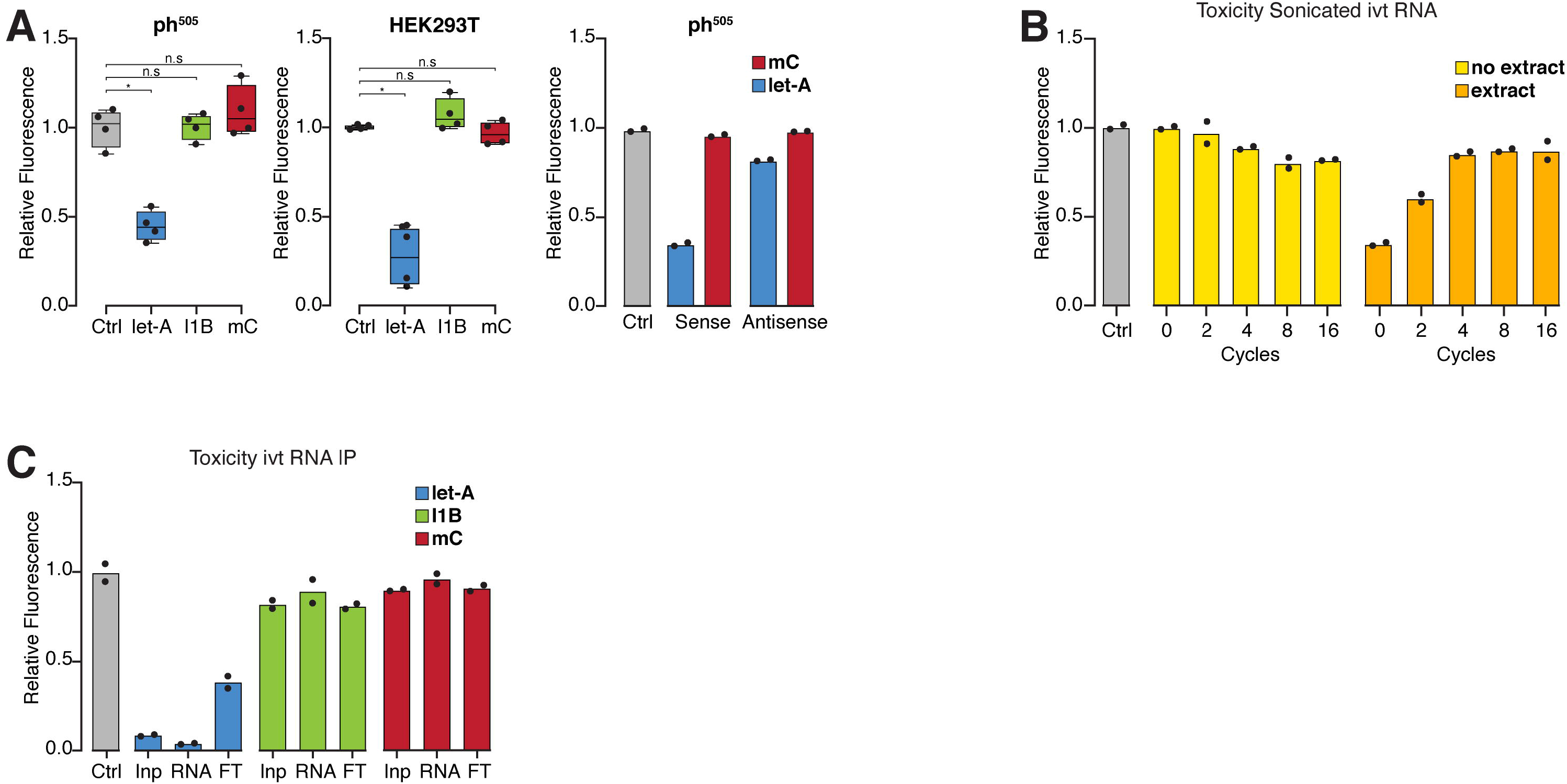
*In vitro* transcribed *let-A* shows an oncolytic effect if incubated in cell extract. (**A**) *In vitro* transcribed (ivt) RNAs (*let-A*, I1B, and mCherry (mC)) were first incubated with *Drosophila* cell extracts and subsequently added to either *ph^505^* cells or HEK293T cells. Only incubated *let-A* RNA showed an oncolytic effect for both *ph^505^* cells and HEK293T cells as quantified by alamarBlue. Neither the I1B and mC sequences, nor the *let-A* antisense sequences had a comparable effect. Bars across conditions denote Wilcoxon Rank-Sum Test p-values (n.s.: non-significant, *: ≤ 0.05). (**B**) Cell viability of *ph^505^* cells treated with sonicated ivt *let-A* RNA. The RNA not incubated with fly cell extract did not show toxicity, but RNA incubated with fly cell extract lost its toxicity with increasing number of sonication cycles. (**C**) Cell viability of *ph^505^* cells treated with affinity purified *in vitro* transcribed *let-A*, *I1B*, or mC RNA. RNAs were transcribed with biotinylated ribonucleotides, incubated with cell extracts, and purified by affinity purification before application to the cells. Only the purified ivt *let-A* RNA showed toxicity. Input, without purification; RNA, affinity purified RNA; FT, flow through after affinity purification.

To further confirm that *let-A* is causally inducing toxicity, we *in vitro* transcribed biotinylated *let-A*, incubated with cell extracts, and isolated the RNA by biotin-streptavidin affinity purification. The purified *let-A* RNA showed stronger toxic effect on *ph^505^* cells compared to the input RNA (Fig. 2C). In contrast, neither *I1B* nor mCherry RNA that were *in vitro* transcribed and processed in the same manner were able to kill the *ph^505^* culture cells (Fig. 2C). We performed total RNA-seq with the affinity purified RNA. As shown in supplementary figure 1B, after affinity purification *let-A* sequence was substantially enriched compared to the input sample. Together, these results strongly suggest that *let-A* RNA is indeed the molecule that induces cell death in *ph^505^* cells. To exert this toxicity, it needs to be modified and/or processed to become active, however.

### *let-A* RNA full length sequences are required to induce cell toxicity

As *let-A* has a length of ~6 kb and needs further modifications and/or processing to become active, we next tried to identify essential regions or short sequences within the full-length transcript. We first generated a series of deletion constructs starting from the 3’ end of *let-A*, and transduced *ph^505^* cells with viral vectors expressing every single construct (Fig. 3A, r1-r16). In contrast to *let-A* full length construct, all the deletion constructs showed a gradually reduced toxicity and resulted in cell death only three days after induction (Fig. 3B). From the RNA-seq analysis (supplementary figure 1B), we observed several regions within *let-A* where short sequencing reads were enriched (Fig. 3A, IVT *let-A* IP sequencing). Based on these sequencing results, we next split full length *let-A* into three shorter fragments (*let-A* p1-p3) (Fig. 3A). Again, when these partial *let-A* fragments were expressed individually in *ph^505^* cells, all three shorter RNAs showed reduced toxicity compared to full length *let-A* (Fig. 3C). Only when two or all three shorter RNAs were expressed together, *ph^505^* cells could be killed as efficiently as full length *let-A* (Fig. 3C). We also tested constructs that only express single short fragment from the most enriched regions in the sequencing results (Fig. 3A). When induced in *ph^505^* cells, all these short fragments showed significantly reduced killing activity to cells (Fig. 3D). We finally used deletion constructs as well as mutagenized *let-A* constructs to identify potential active regions of the lncRNA (Fig. 3E, see supplementary figure 1C for details). For the 80% *let-A* construct (letA80) 20% of *let-A* was removed from the 3’ end by PCR. For letA80-H, part of HygR and for letA80-M, part of mCherry was fused to letA80 on the 3’end to keep transcript size at around 6 kb. For the mutagenized let-A variants (letAmut7 and letAmut13), mutagenesis was performed manually in a semi-random fashion. Briefly, *let-A* was divided into three fragments (*let-A* 5’ 1-1913, 1914-3860, 3861-5947) and bases were manually semi-randomly exchanged while respecting local intramolecular base pairings. While mutagenized *let-A* constructs showed a partially increased cell viability (Fig. 3E), no single region could be identified to be responsible for the overall toxicity. Together these experiments suggest that multiple regions within *let-A* have an additive effect towards toxicity, and the full-length sequence is required to act at maximum efficiency.

**Figure 3.**
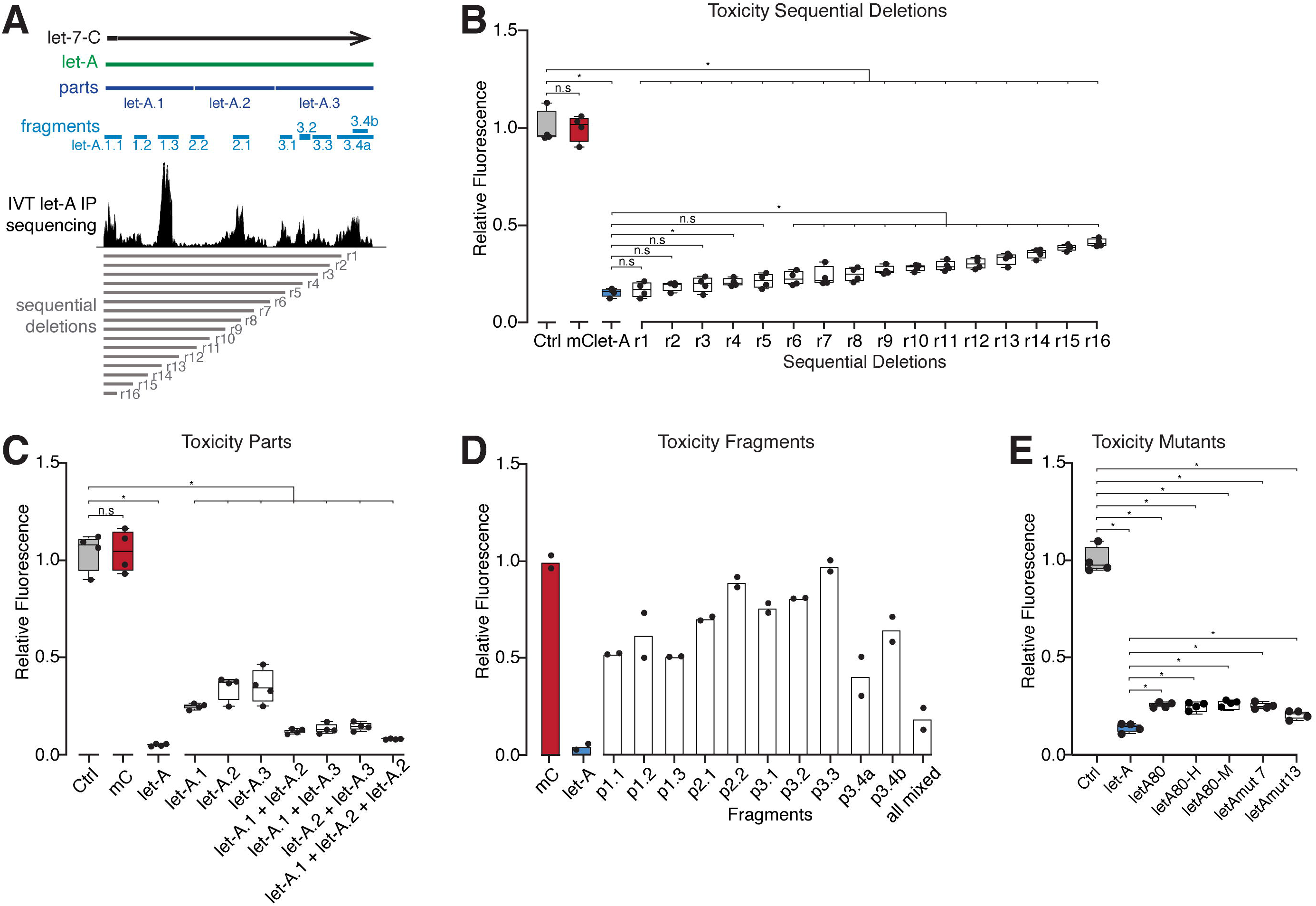
Mapping active sequences in the *let-A* RNA. (**A**) A map showing various constructs for shorter parts, fragments, and sequential deletions within *let-A* sequence. IVT *let-A* IP shows the sequencing reads enriched within *let-A*, the enriched regions were used to design the fragment constructs. (**B**) Cell viability of *ph^505^* cells transfected with constructs encoding a series of *let-A* deletion sequences. Bars across conditions denote Wilcoxon Rank-Sum Test p-values (n.s.: non-significant, *: ≤ 0.05). (**C**) Cell viability of *ph^505^* cells transfected with vectors encoding *let-A* parts p1-p3 as well as combinations thereof. Bars across conditions denote Wilcoxon Rank-Sum Test p-values (n.s.: non-significant, *: ≤ 0.05). (**D**) Cell viability of *ph^505^* cells transfected with vectors encoding *let-A* short fragments depicted in (A). (**E**) Cell viability of *ph^505^* cells transfected with mutagenized *let-A* constructs (see Suppl. Figure 1A). Bars across conditions denote Wilcoxon Rank-Sum Test p-values (n.s.: non-significant, *: ≤ 0.05).

### *In vitro* transcribed *let-A* RNA enters cells

To understand how *let-A* RNA acts to induce cell toxicity, we prepared ivt *let-A* RNA using fluorophore-labeled nucleotides (named *fluo-let-A*, see M&M) for imaging analysis. After incubation with *Drosophila* cell extracts and purification, ivt *fluo-let-A* can induce cell death in both *ph^505^* cells and HEK293T cells, indicating this labelled *let-A* RNA maintains toxicity, however, slightly weaker compared to the ivt RNA with normal nucleotides (supplementary figure 1D). Purified *fluo-let-A* was either incubated with or without cell extracts, and *Drosophila ph^505^* culture cells treated for 1 hour. Image analysis showed fluorescent signals localized both inside and outside of the cells (Fig. 4). After 3D reconstruction of the images, the signals were found to locate both in the cytoplasm and outside of the cells (Fig. 4A, inlet). In contrast, for cells treated by RNA not incubated with cell extract, the fluorescence was only observed outside cells (Fig. 4B, inlet). Next, we treated HEK293T cells with ivt fluo-let-A. Similarly, the fluorescent signals inside HEK293T cells were only observed when the RNA was incubated with cell extract (Fig. 4C, D). These results show that ivt RNA can enter the cells to exerts its function. However, to gain the ability to enter the cells and become active, the RNA needs to be modified/processed.

**Figure 4.**
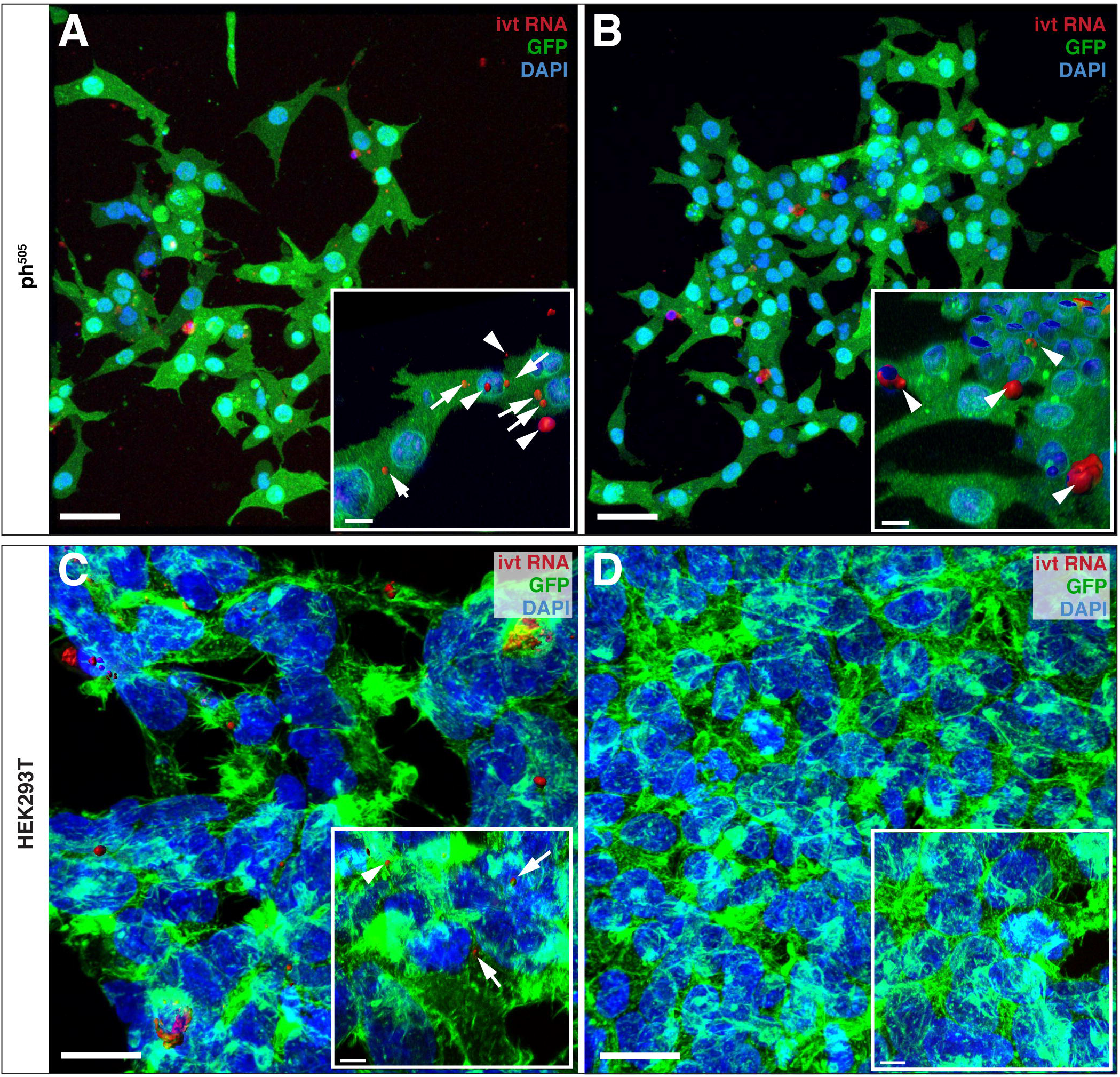
Fluorescently labelled ivt *let-A* RNA enters the cells. (**A**) *let-A* was *in vitro* transcribed using fluorescent nucleotides (fluo-let-A). After incubation with fly cell extract, *ph^505^* cells were treated and fluorescence were imaged by confocal microscopy. Inlets show 3D reconstructions of microscope data. A substantial amount of fluorescence can be observed inside (arrows) as well as outside (arrow head) the cells. (**B**) Without treatment with fly cell extract, fluorescence can only be observed outside of cells (arrow head). (**C**) A similar experiment shows that activated fluo-let-A can also enter HEK293T cells. (**D**) Without the incubation fluorescence can only be observed outside of cells. Scale bars are 20 μm in the main figures, and 5 μm for the inlets.

### Extracellular vehicles and exo/endocytosis are required for *let-A* mediated cellular toxicity

To investigate the mechanisms of *let-A* mediated cellular toxicity, we performed an inhibitor screen using chemicals used to target major components of different cell signaling pathways, exo/endocytosis pathways, cell cycle, as well as cell death. Among these, chemicals used as exo/endocytosis inhibitors showed the strongest effects by increasing cell viability when added to the culture medium after *let-A* induction. Extracellular vehicles (EVs) are capable of transporting RNA cargos and mediating intracellular signaling (Colombo et al., 2014). As such, we next tested if such nanoparticles are possibly required for *let-A* mediated cell toxicity. GW4869 is a neutral sphingomyelinase inhibitor and a currently widely used pharmacological agent for blocking exosome production (Kosaka et al., 2010). Indeed, adding GW4869 to the medium could efficiently decrease the toxicity of *let-A* in *ph^505^* culture cells (Fig. 5A). Similarly, Brefeldin A (BFA), an inhibitor of protein transport from the endoplasmic reticulum to the Golgi complex also reduced *let-A* toxicity increasing cell viability (Fig. 5A). Moreover, we used Dynasore, an inhibitor of dynamin that is required for clathrin-mediated and caveolin-dependent endocytosis (Macia et al., 2006). Dynasore acts to suppress exosome internalization, resulting in a moderate increase in *ph^505^* culture cell viability when *let-A* was induced (Fig. 5A). The effect of all three chemicals appeared to be specific for *let-A*, as cells are not affected when control mCherry RNA was induced (Fig. 5A). Furthermore, adding GW4869 or BFA into the culture also increased the cell viability of HEK293T cells when *let-A* was induced in these cells, whereas BFA has no effect (Fig. 5A). These experiments show that cellular processes like exocytosis and endocytosis are required for *let-A* induced cell toxicity both in *ph^505^* culture as well as HEK293T cells.

**Figure 5.**
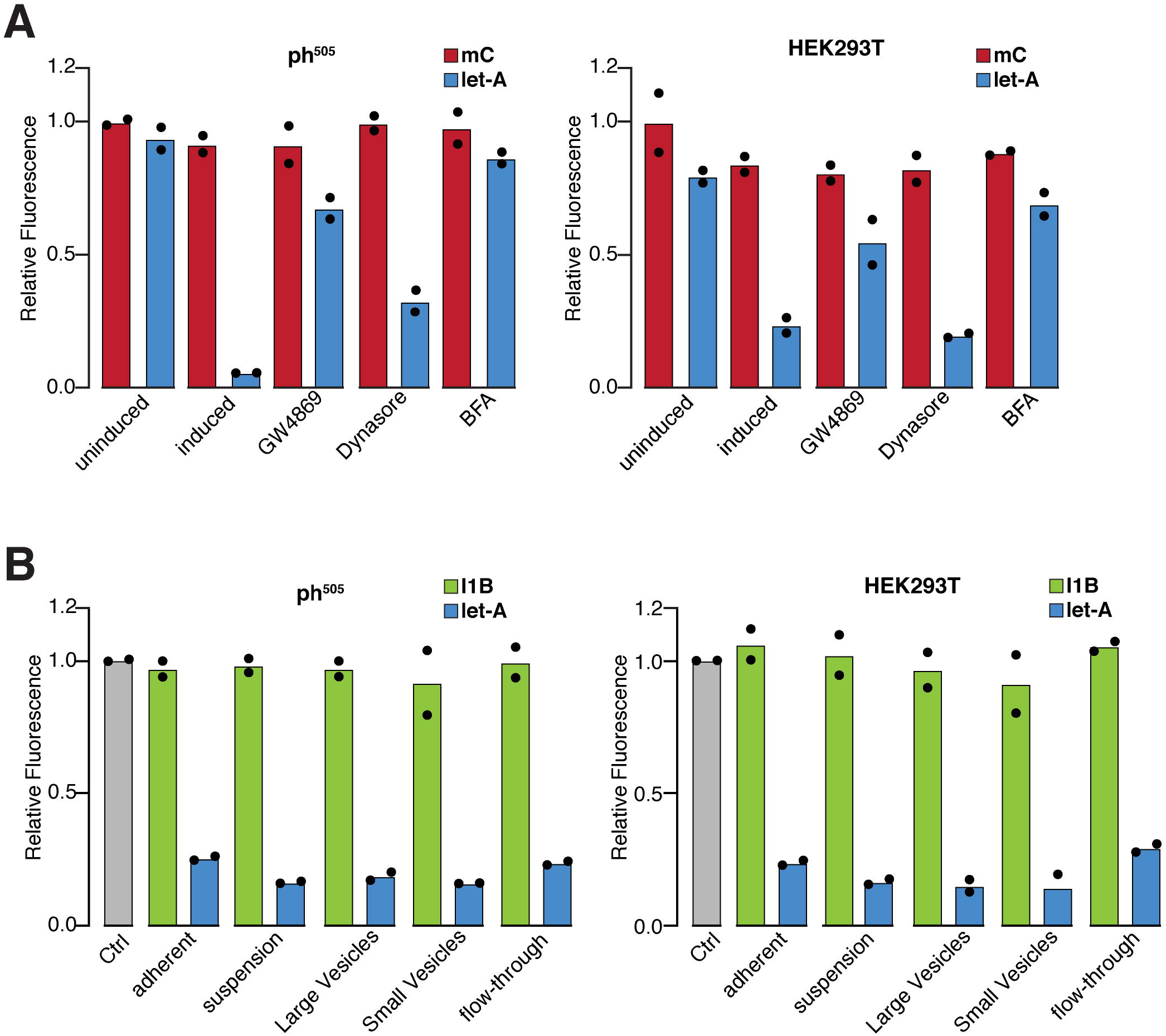
Extracellular vehicles and exo/endocytosis are required for *let-A* mediated cellular toxicity. (**A**) *ph^505^* cells as well as HEK293T cells were treated with conditioned let-A/medium in the presence of inhibitors of exo- or endocytosis. In most cases toxicity was substantially reduced demonstrating the need for packaging *let-A* in an extracellular vehicle like exosomes and the uptake of these vehicles via endocytosis. Inhibitors had no effect on mCherry control constructs. (**B**) Exosomes were collected from conditioned let-A/medium and used to treat either *ph^505^* or HEK293T cells. Exosome fractions from let-A/medium retained toxicity but control fractions from I1B/medium not.

Next, we tried to collect exosomes from *ph^505^* cells expressing *let-A* or I1B by centrifugation and column purification using an exosome collection kit and test their toxicity. While RNA from all the fractions from I1B-expressing cells has no effect on *ph^505^* cells, RNA from all *let-A* fractions was toxic (Fig. 5B).

### Assessing cellular signaling pathways responding to *let-A* treatment

To further uncover the underlying death mechanism in both *Drosophila* and mammalian cells, we investigated what signaling pathways might be involved. Based on the readout of the inhibitor screen mentioned above, we focused on Toll signaling pathway. Toll-like receptors (TLRs) are a family of membrane proteins that can induce an immune response upon recognition of microbial pathogens. First discovered in *Drosophila*, this family of proteins is evolutionarily conserved from flies to humans. In particular, several TLRs (TLR3, TLR7, TLR8, and TLR9) are known to recognize either double stranded or single stranded RNAs to initiate downstream signaling events (Alexopoulou et al., 2001). To test if the Toll signaling pathway is required for the *let-A* induced cell death, we applied several inhibitors that targeted different components in *ph^505^* cells (Fig. 6A). When receptor TLR3 was blocked, an increase in cell viability was observed (Fig. 6A). When inhibitors against more downstream components were applied, including the adaptor protein MyD88 and the kinase TBK1, we observed a further increase in cell viability (Fig. 6A). In addition, we used two inhibitors for the most downstream transcription factor, NF-κB, which also resulted in a better cell survival (Fig. 6A). The same inhibitors were used to block Toll signaling in mammalian HEK293T cells. Similarly, treatment could inhibit *let-A* induced toxicity in the HEK cells and increasing significantly cell viability (Fig. 6B).

**Figure 6.**
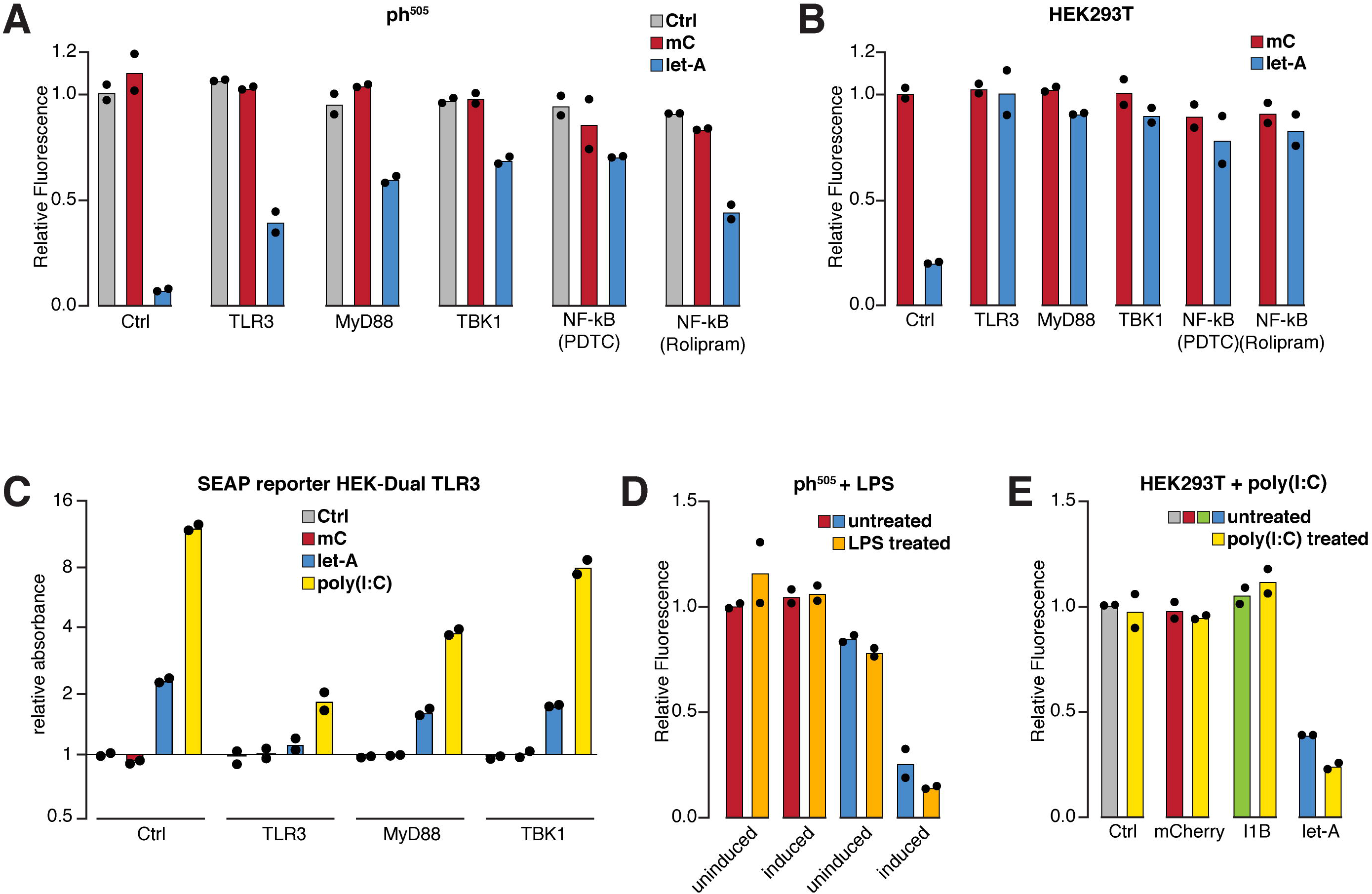
Toll pathway reacts to *let-A* treatment. (**A**) Cell viability of *ph^505^* cells treated with inhibitors against Toll signaling components TLR3, MyD88, and TBK1, as well as with two NF-κB inhibitors. Cells were first incubated with different inhibitors for one hour, then treated with purified RNAs from mCherry/medium or let-A/medium for three hours. (**B**) Cell viability of HEK393T cells treated with inhibitors against Toll signaling components. Cells were treated by the same inhibitors and processes as (**A**). (**C**) SEAP activity in HEK-Dual hTLR3 cells treated with purified RNA from let-A/medium, mCherry/medium, or poly(I:C), with or without Toll signaling inhibitors. (**D**) Cell viability of *ph^505^* cells with or without pre-treatment with LPS before *let-A* (blue) or mCherry (red) expression was induced. Note that *let-A* induction could kill the LPS pre-treated cells even faster. (**E**) Cell viability of HEK393T cells with or without pre-treated with poly(I:C), before treated with purified RNA from mCherry/medium or let-A/medium. let-A/medium purified RNA could kill the poly(I:C) pre-treated cells much faster.

We next used a HEK-Dual hTLR3 cell line expressing SEAP (Secreted embryonic alkaline phosphatase) as a reporter for Toll signaling. The SEAP activity was low in the untreated condition, but substantially increased upon treatment with poly(I:C) (polyinosine-polycytidylic acid), a known inducer of the Toll signaling pathway (Fig. 6C). When HEK-Dual hTLR3 cells were treated with mCherry/medium, we did not observe an increase in SEAP activity (Fig. 6C). Conversely, treating cells with let-A/medium increased SEAP activity (Fig. 6C), indicating induction of Toll signaling. Furthermore, when Toll pathway inhibitors were added, a strong reduced of SEAP activity in both poly(I:C) treated and *let-A* treated cells was observed (Fig. 6C). Lastly, we pre-treated *ph^505^* cells with LPS (lipopolysaccharide), which is an inducer of Toll signaling but does not cause cell death (Fig. 6D), to sensitize the cells to other activators of Toll signaling. When mCherry was induced in LPS pre-treated *ph^505^* cells, no effect was observed (Fig. 6D). In contrast, when *let-A* was induced in LPS pre-treated *ph^505^* cells, this killed cells much faster and all cells died within one hour (Fig. 6D). Similarly, poly(I:C) pre-treated HEK cells also became more sensitive to *let-A* and showed a further reduced viability compared to cells untreated with poly(I:C) (Fig. 6E). Together, these results show that Toll signaling pathway is required for *let-A* induced cell death in both *Drosophila* and human cancer cells.

It is well known that dsRNA can induce an interferon response in mammalian cells (Gantier & Williams, 2007). In contrast to the classical inducer poly(I:C), no activation of Interferon ß is observed after exposure of HEK293T cells to *let-A* (supplementary figure 1E), however. Levels of induction by *let-A* are comparable to control RNA (mCherry) or the fragment I1B. This suggests that we do not observe a general activation of the innate immune response of mammalian cells to foreign RNA but a rather specific activation of the Toll pathway, leading to cell death.

### Reduction of nucleolus size as a consequence of *let-A* treatment

We have shown previously, that the ecdysone pulse at metamorphosis of *D. melanogaster* triggers expression of the *let-7-Complex*, transforming the tumorigenic *ph^505^* cells in the imaginal discs into “metamorphed” non-tumorigenic state (Birbaumer et al., 2021; Jiang et al., 2018). In order to assess the cellular events following transcription of *let-7-C* we treated cultured *ph^505^* cells with ecdysone and analyzed transcription patterns by RNA-seq over time. Among a number of pathway changes, we observed that after a 24-hour ecdysone treatment the GO term “ribosome biogenesis” tops the list. Several ribosome-associated genes, like pol I components or ribosomal protein genes, were found down-regulated. Indeed, by staining nucleoli with fibrillarin we observed a visible nucleolus reduction in metamorphed cells compared to the tumorigenic *ph^505^* cells. There is a well-accepted correlation between increased ribosome biogenesis, abnormal nucleoli and cancer (Ma et al., 2016; Núñez Villacís et al., 2018). We now observe a similar reduction if *ph^505^* cells are treated with let-A medium alone (Fig. 7A). This suggests that *let-A* is one of the components of *let-7-C* triggering the downregulation of genes involved in ribosome biosynthesis. A similar effect is observed if HEK293T cells are treated with let-A medium (Fig.7B). Given the multiplicity of nucleoli observed in mammalian cells, after *let-A* treatment we observe a stronger clustering indicated by a reduced overall number of nucleoli. These results suggest that *let-A* exposure triggers a defined cellular response by downregulating components of the translational apparatus.

**Figure 7.**
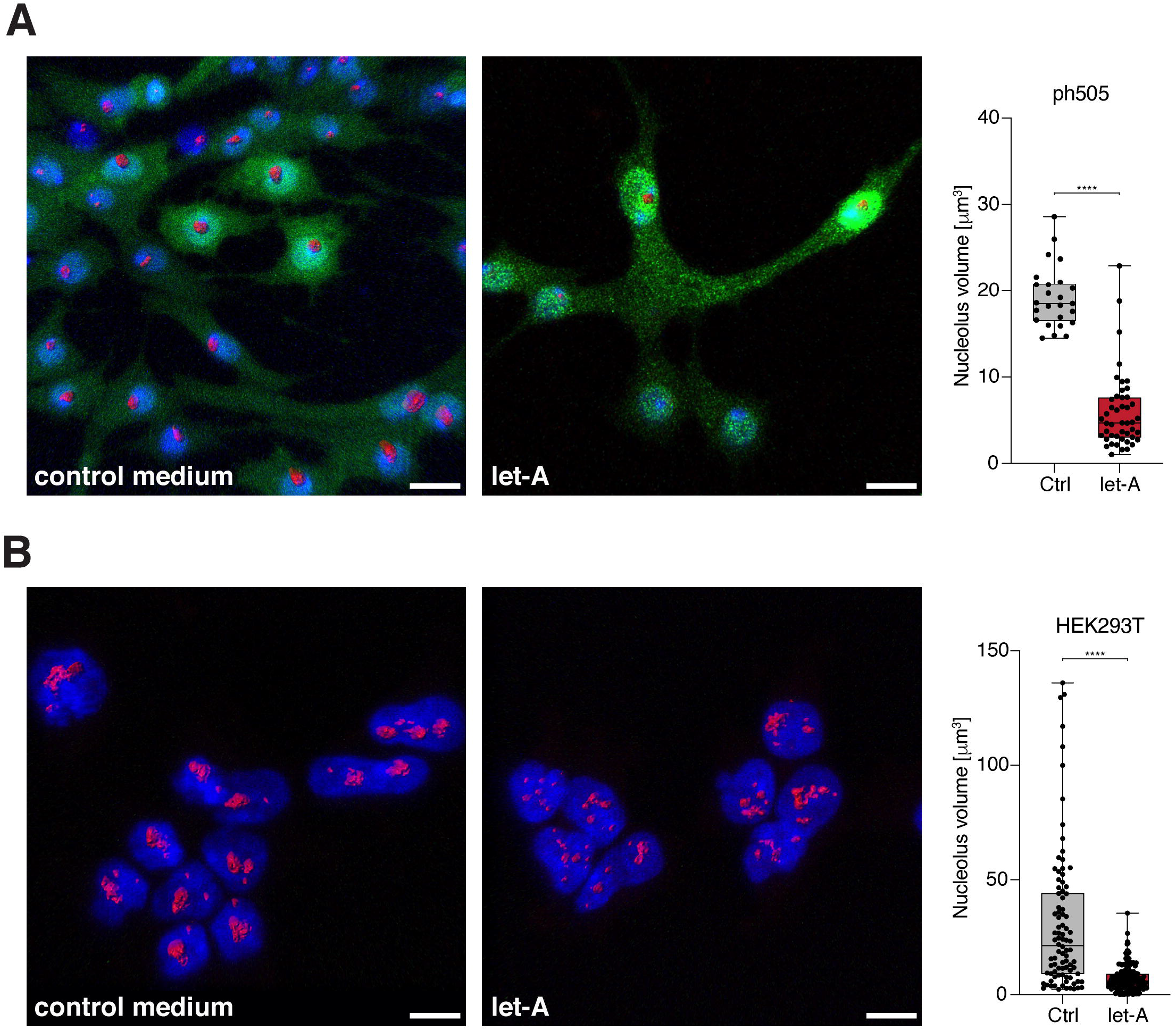
*let-A* reduces the nucleolus’s volume in treated cells. (**A**) Confocal images showing *ph^505^* cells treated with mCherry/medium (control) or let-A/medium (let-A). Immunostaining using Fibrillarin antibody (nucleolus, red). Right panel: Quantification of the nucleolus volume showing that the nucleolus volume was reduced in let-A/medium treated *ph^505^* cells. Bar across conditions denotes Wilcoxon Rank-Sum Test p-values (****: ≤ 0.0001). (**B**) Confocal images showing HEK293T cells treated with mCherry/medium (control) or let-A/medium (let-A). Immunostaining using Fibrillarin antibody (nucleolus, red) and DAPI for nucleus (blue). In mCh/medium treated cells, each nucleus contains one large Fibrillarin staining (volume larger than 25 μm^3^) and a few smaller ones. But in let-A/medium treated cells, the larger structure disappears and more smaller structures are observed in the nucleolus. Right panel: Quantification of the volume of nucleoli in mCh or let-A/medium treated HEK293T cells, showing that the nucleolus volume was reduced in let-A/medium treated cells. Bar across conditions denotes Wilcoxon Rank-Sum Test p-values (****: ≤ 0.0001). Scale bars in all panels are 10 μm.

## Discussion

In our accompanying paper (Birbaumer et al., 2021), we report the discovery of *let-A*, a long non-coding RNA (lncRNA) in the *Drosophila let-7* complex and show that *let-A* can induce cell death in *Drosophila* cancer cell culture and suppress the growth of tumors *in vivo*. In this study, we further show that *let-A* can also induce oncolytic death in mammalian cancer cells, therefore exhibit an evolutionary conserved anti-cancer function. We show that either induced expression of *let-A* or adding RNA purified from the *Drosophila let-A* conditioned medium results in HEK293T cell death similarly as we have shown with *Drosophila ph^505^* cells. HEK293T is an immortalized cell line originating from embryonic kidney, which can form tumors in mouse models thus making it a widely used cell culture in cancer research. Moreover, we find that *let-A* is toxic to a wide range of different human cancer cells, including human cervical cancer (HeLa), human breast cancer (Mcf7), human glioblastoma (BT8A), human leukemia (K562), and human pancreatic cancer (Mia PaCa-2). On the contrary, the toxic effect is not observed with non-tumorigenic and more differentiated cell lines, including human fibroblast cells (HS68), human primary mesenchymal stem cells (MSC), and differentiated adipocyte and smooth muscle cells. Therefore, *let-A* appears to have a toxic specificity towards cancer/immortal cells but not cells derived from normal tissue, which is an obvious benefit for a potentially novel cancer therapeutic RNA. Such a specific effect is best shown when HEK293T and HS68 cells are co-cultured and treated with *let-A*, which results in death of HEK293T cells but not HS68 cells. This specificity can be explained either by a mechanism that is only active in cancer cells, or by an inhibitory mechanism presents in normal cells. To understand this, an un-biased screening for relevant pathways might provide insights into *let-A*’s cancer cell specificity.

Unlike some well-studied lncRNAs, *let-A* is much longer and has a sequence of 6 kb. We performed intensive analyses to try to characterize the most essential regions within the *let-A* RNA. So far, we have not been able to identify single short regions in *let-A* that would show the same efficiency of toxicity as the full-length RNA. However, shorter fragments representing different regions of *let-A* show an additive effect. This suggests that different regions within *let-A* are needed to act together to mediate the toxic effect. Full length *in vitro* transcribed (ivt) *let-A* RNA is not toxic by itself but becomes active after incubation with cell extracts, indicating cellular modification and processing are necessary to generate the active fragments. RNA-seq analysis shows the enrichment of sequences from several parts in *let-A*, suggesting candidate regions for the active fragments. As potential mechanism *let-A* may be processed into shorter fragments containing siRNAs, miRNAs or alternative short RNAs with specific secondary structures to exert their functions in the target cells. Alternatively, the sequences may also act as RNA sponges to inhibit the function of other RNA molecules within the cells.

We observe that fluorophore-labeled *let-A* RNA can enter the cells, but only when the RNA is pre-incubated with cell extracts. Interestingly, also resistant cells appear to take-up the labeled RNA suggesting that the toxicity pathway is triggered after entry into the cell. We also provide evidence that the processed *let-A* RNA might be carried by the EVs to act on neighboring or distant cells. By treating cells with chemicals that are known to block exocytosis or endocytosis, the toxicity induced by the expression of *let-A* can largely be suppressed. In addition, enrichment of *let-A* RNA can be detected in the purified cellular fraction containing EVs. EVs are cell derived nanostructures with enclosed membrane, which have been shown to act as extracellular transporters between distant cells and mediate intracellular communication (Pegtel & Gould, 2019). Among the various types of cargos that EVs carry, a large subset are different RNA species (O’Brien et al., 2020). During the past decade, EVs have shown promise for clinical applications as the next generation therapeutic delivery vehicles (Luan et al., 2017). Hence, the association of *let-A* with EVs further endorses the view that this lncRNA has a potential for therapeutic applications.

We show *let-A* may exert its toxicity via evolutionarily conserved Toll signaling pathway. Members of the Toll-like receptor family (TLR3, TLR7, TLR8, and TLR9) are known to recognize either double stranded or single stranded RNAs to initiate downstream signaling (Alexopoulou et al., 2001). One of the observable consequences of *let-A* treatment at the cellular level is the reduction of nucleolar size. This has intriguing ramifications as nucleolar size and in consequence translational efficiency is coupled to tumorigenic performance (Núñez Villacís et al., 2018). Taken together, we propose a model for the oncolytic effect mediated by *let-A* RNA: first, *let-A* is induced in particular cells; second, the full-length RNA is modified and processed; third, the processed RNAs are released and transported in EVs and therefore mediating the toxicity in other cells; last, the active fragments of *let-A* are recognized by TLRs initiating downstream processes like nucleolar shrinkage and eventually cell death.

The 2D tumor cell cultures used in labs for decades are known to show disadvantages in many aspects compared to cancers grown *in vivo*. Conversely, 3D organoid models have become a powerful tool to improve physiological comparability (Driehuis et al., 2019; Huang et al., 2015). Indeed, we have preliminary evidence that also in an organoid model using tumor tissue collected from pancreatic ductal adenocarcinoma (PDAC) patients *let-A* diminishes the growth of this cancer organoid (unpublished results). In summary, the oncolytic activity of *let-A* across species, the broad toxicity in various cancer types, the specificity towards cancer cells, and the efficiency shown in human cancer tissue together these findings make *let-A* a promising RNA for a new biomolecule-driven treatment of cancer.

---

**Supplementary Figure 1**

(**A**) Toxicity as measured by alamarBlue of ivt RNA on *ph^505^* cells after incubation in different conditions or with different cell extracts. Left panel: ivt RNA was either directly added to cells (ivt only) or first incubated with fly cell extracts with ATP (extract) and without (extract – ATP), M3 medium with ATP (medium +ATP) and extracts pretreated with proteinase K or cyanase (extract +proteinase K and extract +cyanase, respectively). Right panel: ivt RNA was incubated with extracts of four different cell lines. (**B**) RNA-seq of the input or affinity purified *in vitro* transcribed RNA as in (Fig. 2C). Left: Number of reads per kilobase and million reads (RPKM) showed the enrichment of *let-A* after purification. Right: Top 20 genes with most counts in the affinity purified sample and their log2 fold changes compared to the input sample. Let-7-C was the most enriched RNA. (**C**) Comparison of cellular toxicity by mutagenized variants of *let-A*. 20% of *let-A* is removed from the 3’-end (let-A80) or replaced by a neutral sequence (part of hygromycin; let-A80-H or part of mCherry; let-A80-M). let-Amut 7 and let-Amut 13 represent two variants containing approx. 20% base exchanges over the entire sequence of *let-A*. white box: consensus *let-A* sequence, black box: mutagenized sequences compared to consensus *let-A*, line: deleted sequence. Toxicity was tested in Fig. 3E. (**D**) Fluorescently labelled *let-A* RNA (fluo-let-A) shows a similar toxicity as regular RNA (let-A). (**E**) Assessing an interferon response of HEK293T cells after RNA exposure. HEK293-TLR3 cells were exposed to *let-A* or I1B RNAs. Poly(I:C) was used as a standard inducer of the interferon response. INFß is not induced by *let-A*. Comparably, TNF is moderately induced, while more downstream components like CXCL2 and NFkB are not induced. Bars across conditions denote Wilcoxon Rank-Sum Test p-values (n.s.: non-significant, *: ≤ 0.05).

## Materials and methods

### *ph^505^* cell line generation

For establish *ph^505^* cell line, two to three 1 week old transplanted *ph^505^* tumors were isolated from the host flies and dissected into small pieces in sterilized PBS following 3 times wash with Shields and Sang M3 Insect medium (US Biological) containing Penicillin/streptomycin(P/S). The cells were resuspended in 1ml of M3 (P/S) medium and plated into the 96 well plate 200μl per well. After the stable cell growth, single cell line was isolated using Terasaki plate (Greiner Bio-One). To confirm the cell line, genomic DNA was isolated and performed sequence analysis.

### Cell culture and treatments

*Drosophila ph^505^* cell lines were maintained at 25 °C in Shields and Sang M3 Insect medium (US Biological), supplemented with 2% Fetal Bovine Serum (FBS) (PAN Biotech), 2.5% fly extract, 5◻μg/ml human insulin (Sigma), 100 U/ml penicillin and 100◻μg/ml streptomycin (Gibco, Life Technologies). S2 cells were cultured in Schneider’s medium (Gibco, Life Technologies) supplemented with 10% FBS (PAN Biotech) at 25 °C. *Spodoptera frugiperda* Sf21 cells (Clontech) were propagated in Grace’s medium (Gibco, Life Technologies) with 10% FBS (PAN Biotech) at 27°C. All mammalian cell lines except for the mesenchymal stem cells were grown in Dulbecco’s Modified Eagle’s Medium (DMEM) high glucose (Sigma) supplemented with 10% FBS (Sigma) and 2 mM L-glutamine (Gibco). For HEK-Dual hTLR3 cells (InvivoGen) the growth medium was additionally supplemented with 100 μg/ml Normocin, 100 μg/ml Hygromycin B and 50 μg/ml Zeocin (all from InvivoGen). Primary human bone marrow-derived mesenchymal stem cells were obtained from ATCC (PCS-500-012). They were maintained according to manufacturer’s instructions and differentiated as described in (Almalki & Agrawal, 2016).

HEK293T cells were transfected with using Fugene HD (Promega) following manufacturer’s instructions. After two days transcription was induced with 1 μg/ml tetracycline (Sigma) and cell viability was measured after overnight incubation. SEAP activity was detected using QuantiBlue (InvivoGen) and the absorbance quantified on a Tecan Infinite M1000 PRO microplate reader at 650 nm. The following inhibitors were used: TLR3/dsRNA Complex Inhibitor (Sigma) 30 μM; MyD88 inhibitor T6167923 (Anawa) 50 μM; TBK1 inhibitor MRT67307 (Sigma) 2 μM; and the NF-kB inhibitors Rolipram (Abcam) 1 μM; pyrrolidine dithiocarbonate (PDTC, Sigma) 25 μM. For all the inhibitor experiments, cells were pretreated one hour with the respective inhibitor before induction or addition of the extracted medium. For induction, the cell viability was determined after three hours, for extracted medium treatment after one hour, for *Drosophila* cell lines and for mammalian cell lines after overnight incubation.

For activation of the Toll pathway cells were pretreated for one hour with 10 μg/ml LPS (Sigma) or 5 μg/ml poly(I:C) HMW (InvivoGen). Two hours after induction cell viability was measured with alamarBlue. To analyze the interferon response HEK-Dual hTLR3 cells were treated with 5 μg/ml poly(I:C) HMW (InvivoGen) or the different extracted media for four hours, then total RNA was extracted using Trizol (Invitrogen) and Direct-zol MiniPrep kit (Zymo Research) and qPCR was performed as described below.

### Generation of recombinant baculoviruses

Cloning was performed following the strategy described in (Lee et al., 2000). The *let-A* and *I1B* fragments were amplified from S2 cell genomic DNA using the Expand Long Template PCR System (Roche) and cloned in pBacPAK8_EGFP under a metallothionein promoter. *let-B* was cloned from pBacPAK8_EGFP_I1B. Viruses were generated, amplified, analyzed and harvested using the BacPAK Baculovirus Expression System (Clonetech) according to manufacturer’s instruction.

### Cloning for mammalian cell expression

For inducible expression in mammalian cells, *let-A*, *I1B*, mCherry and EGFP were cloned from pBacPAK8 into pSBtet (Addgene) under the control of an inducible TRE promoter. For *in vitro* transcription the fragments were cloned in pGEM-T Easy plasmids under the control of a T7 promoter.

### Cell viability measurement

Cell viability was measured after the indicated time using alamarBlue cell viability assay reagent (Thermo Fisher Scientific) following the manufacturer’s instructions. Fluorescence at 650 nm was measured with Tecan Infinite M1000 PRO microplate reader.

### RNA extraction, cDNA synthesis and Quantitative real-time PCR

RNA from cells or medium was extracted using TRIzol (Invitrogen) following manufacturer’s instructions. Reverse transcription was done with the first Strand synthesis kit (Fermentas) using oligo (dT)18 as a primer and qPCR was carried out with LightCycler 96 (Roche) using GAPDH.

### *in vitro* RNA transcription, purification and sequencing

RNA was transcribed from linearized pGEM plasmids using the MEGAscript T7 Transcription kit (Ambion) following the manufacturer’s instructions. The IVT products were purified using the MEGAclear kit (Ambion). Concentration and size of the transcripts were determined using TapeStation 4200 (Agilent).

Active cellular extract was prepared as described (Crevel & Cotterill, 1991) with slight modifications. Cells were washed twice with PBS and then 43*10^6 cells were dounced on ice in a dounce homogenizer (Pestle B) in extraction buffer (10 mM HEPES, pH 7.5, 10 % v/v ethyleneglycol, 250 mM sucrose, 100 mM NaCl, 2.5 mM MgCl2, 1 mM EDTA, 2 mM DTT, 2 mM ATP, PhosSTOP Phosphatase Inhibitor Cocktail (Roche) and cOmplete™ Protease Inhibitor Cocktail (Roche)). The lysate was centrifuged for 10 min at 20’000g at 4°C. The supernatant was taken and total protein concentration was determined by Pierce BCA Protein Assay Kit (Thermo Fisher Scientific). 35 μl of extract (about 4 mg/ml protein) was mixed with 375 ng of IVT RNA (diluted in 15 μl extraction buffer), incubated at 25 °C for 15 minutes and then the RNA was purified up to twice by TRIzol extraction. Cells used for extract generation were mostly ph505 cells, but same results were obtained with HEK293T, Sf21 and NIH-3T3 cells.

Biotinylated RNA was produced by in vitro transcription in the presence of Desthiobiotin-16-UTP (TriLink Biotechnologies). The resulting transcripts were incubated with active cellular extract. To isolate the biotin labelled RNA, the mixture was incubated with 20 μl of Dynabeads MyOne Streptavidin T1 (Invitrogen) for 15 minutes at 25 °C. Afterwards, another 20 μl beads were added and again incubated 15 minutes at 25 °C. Next, the beads were washed twice with extraction buffer and eluted in elution buffer (extraction buffer with 2 mM avidin (IBA)) at 25°C for 15 minutes. The eluates and flow-through were purified by TRIzol extraction.

The purified RNA was sequenced by first depleting ribosomal RNA with the Ribo-Zero Gold rRNA Removal Kit and then subjecting the remaining RNA to the Illumina TruSeq Stranded Library Prep Kit. The resulting libraries were sequenced on a NextSeq500 at paired end 38.

To fluorescently label IVT RNA the ATTO647N RNA labelling kit (Jena Bioscience) was used. 1/20 of total UTP was replaced with Atto 647N UTP for transcription of the fluorescent labelled IVT RNA. The reaction mix was purified by lithium chloride precipitation and concentration and size of the transcripts were measured using TapeStation 4200 (Agilent).

### Immunohistochemistry and antibodies

Cells were cultured for one day on fibronectin coated culture slides (BD). After medium treatment cells were fixed in 4% paraformaldehyde in PBS for 15 minutes at room temperature. Fixation was stopped by adding 100 mM glycine for five minutes. Next, samples were washed twice with PBS, permeabilized with 0.5% Triton X-100/PBS for three minutes and washed again twice with PBS. Cells were incubated with primary antibody for one hour at room temperature, washed three times with PBS, incubated with secondary antibody for one hour at room temperature and washed three times with PBS. Then samples were incubated with DAPI (1:200 in PBS, Sigma) for 10 minutes and mounted using vectashield (Vectorlabs). Primary antibody used in this study was: mouse anti-fibrillarin (1:100; Abcam ab4566); Secondary antibody was: Alexa 647-conjugated anti-mouse (1:500; Molecular Probes).

### Microscopy

Immunofluorescent images were recorded on a Leica TCS SP8 confocal microscope. Images were processed using ImageJ, Imaris, Photoshop, and Adobe Illustrator.

### Bioinformatics analyses

Short reads were aligned to BDGP dm6 genome assembly using TopHat 2.0.12 for Bowtie 2.2.3 with parameters “--very-sensitive” and a segment length of 18 for the IVT reads. From the aligned reads, differential expression was called using R 3.5.1 and edgeR 3.24.3.

## Data availability

Deep sequencing data for in vitro transcription experiments can be found in GEO with accession GSE186536.

## Acknowledgments

This research was supported by ETH Zürich. We thank Martin Fussenegger for hosting TB, YJ and RP in his laboratory and Andreas Hierholzer for comments on the manuscript. Sequencing was performed by the Genomics Facility Basel.

## Competing interests

The authors filed a patent application related to let-A.

## Notes

### Summary of Updates

Data availability: Added GEO accession number

