## Supplementary Figure for "Inducing oncolytic cell death in human cancer cells by the long non-coding RNA *let-A*": Supplementary Figure 1.pdf

(**A**) Toxicity as measured by alamarBlue of ivt RNA on $ph^{505}$ cells after incubation in different conditions or with different cell extracts. Left panel: ivt RNA was either directly added to cells (ivt only) or first incubated with fly cell extracts with ATP (extract) and without (extract – ATP), M3 medium with ATP (medium +ATP) and extracts pretreated with proteinase K or cyanase (extract +proteinase K and extract +cyanase, respectively). Right panel: ivt RNA was incubated with extracts of four different cell lines. (**B**) RNA-seq of the input or affinity purified *in vitro* transcribed RNA as in (Fig. 2C). Left: Number of reads per kilobase and million reads (RPKM) showed the enrichment of *let-A* after purification. Right: Top 20 genes with most counts in the affinity purified sample and their log2 fold changes compared to the input sample. Let-7-C was the most enriched RNA. (**C**) Comparison of cellular toxicity by mutagenized variants of *let-A*. 20% of *let-A* is removed from the 3'-end (let-A80) or replaced by a neutral sequence (part of hygromycin; let-A80-H or part of mCherry; let-A80-M). let-Amut 7 and let-Amut 13 represent two variants containing approx. 20% base exchanges over the entire sequence of *let-A*. white box: consensus *let-A* sequence, black box: mutagenized sequences compared to consensus *let-A*, line: deleted sequence. Toxicity was tested in Fig. 3E. (**D**) Fluorescently labelled *let-A* RNA (fluo-let-A) shows a similar toxicity as regular RNA (let-A). (**E**) Assessing an interferon response of HEK293T cells after RNA exposure. HEK293-TLR3 cells were exposed to *let-A* or I1B RNAs. Poly(I:C) was used as a standard inducer of the interferon response. INFß is not induced by *let-A*. Comparably, TNF is moderately induced, while more downstream components like CXCL2 and NFkB are not induced. Bars across conditions denote Wilcoxon Rank-Sum Test p-values (n.s.: non-significant, *: $\leq 0.05$).

### Supplementary Figure

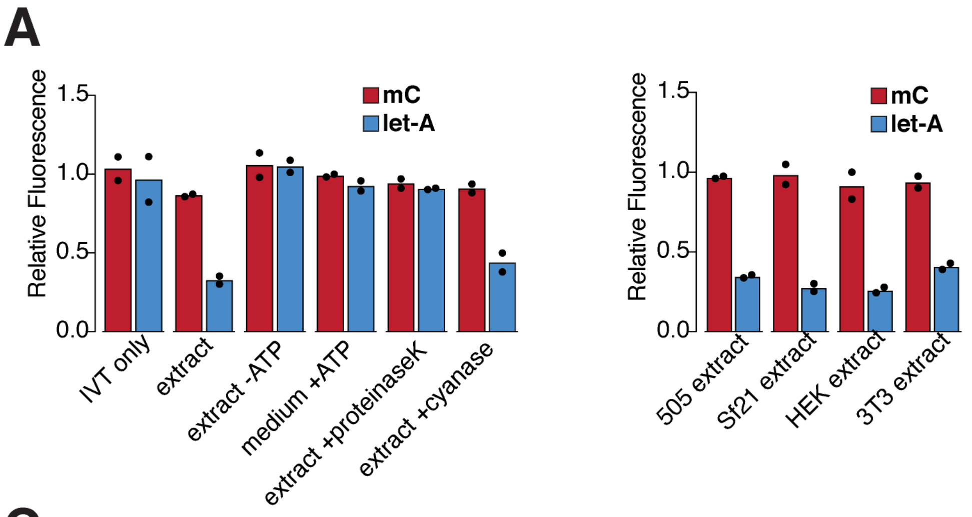


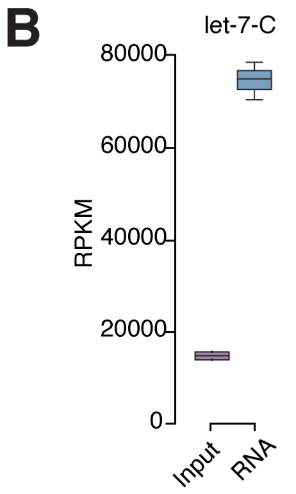


| Gene | Inp | RNA | log2FC |
|---|---|---|---|
| let7C | 13362 | 74352 | 2.48 |
| 28S | 57219 | 31227 | -0.87 |
| 28S | 29377 | 15942 | -0.88 |
| eEF2 | 10808 | 12498 | 0.21 |
| eEF1a1 | 14941 | 11689 | -0.35 |
| Arc1 | 18066 | 8989 | -1.01 |
| CR10102 | 15603 | 7758 | -1.01 |
| 28S | 9041 | 6583 | -0.46 |
| CG10365 | 1432 | 6424 | 2.16 |
| aTub84B | 7345 | 5629 | -0.38 |
| Hsc70-4 | 6257 | 5069 | -0.30 |
| pAbp | 3136 | 4844 | 0.63 |
| Act5C | 5387 | 4842 | -0.15 |
| RpL15 | 5187 | 4705 | -0.14 |
| Hsp83 | 4592 | 4635 | 0.01 |
| RpS3A | 5344 | 4621 | -0.21 |
| 28S | 6981 | 4078 | -0.78 |
| pre-rRNA | 3106 | 3553 | 0.19 |
| RpS29 | 3062 | 3498 | 0.19 |
| RpS23 | 4141 | 3357 | -0.30 |

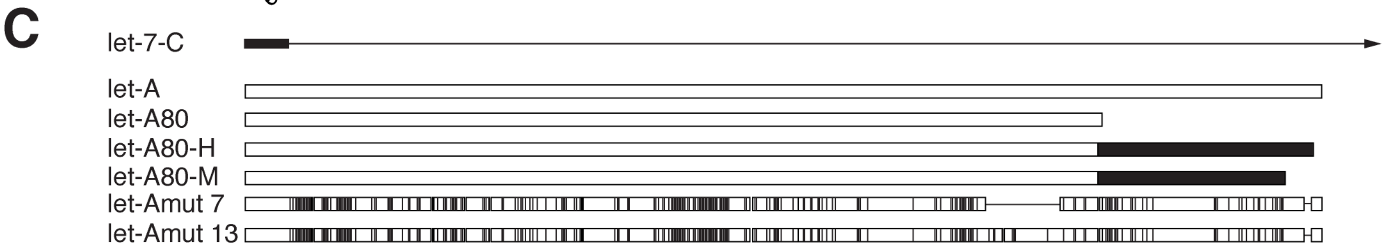


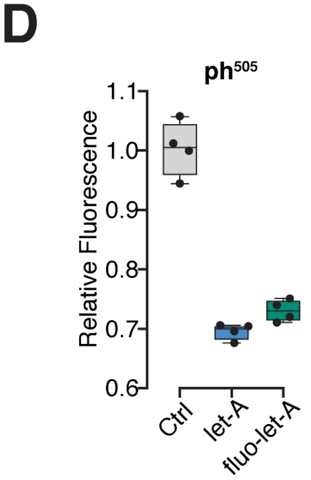


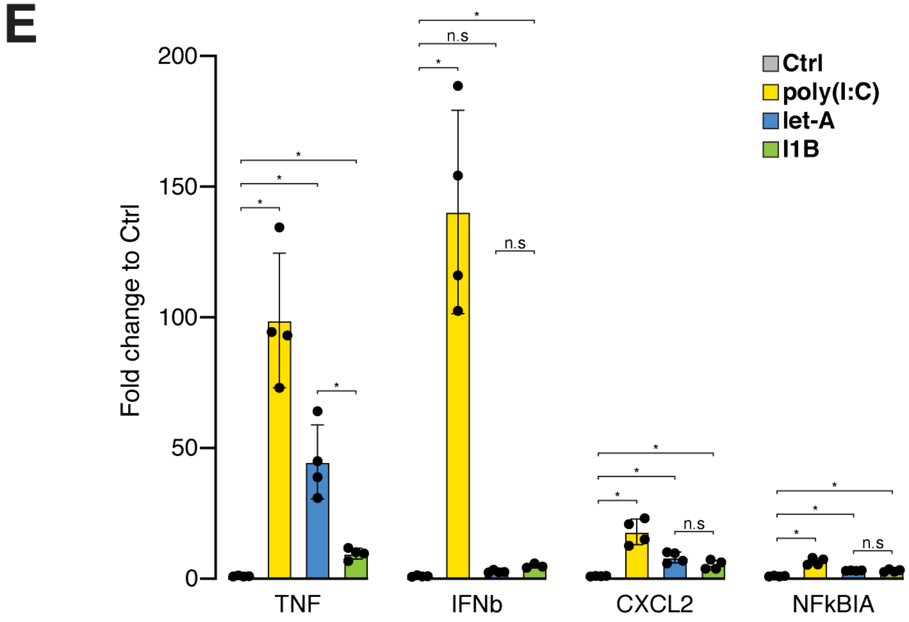
